# Data-driven machine learning models for decoding speech categorization from evoked brain responses

**DOI:** 10.1101/2020.08.03.234997

**Authors:** Md Sultan Mahmud, Mohammed Yeasin, Gavin M. Bidelman

## Abstract

Categorical perception (CP) of audio is critical to understand how the human brain perceives speech sounds despite widespread variability in acoustic properties. Here, we investigated the spatiotemporal characteristics of auditory neural activity that reflects CP for speech (i.e., differentiates phonetic prototypes from ambiguous speech sounds). We recorded high density EEGs as listeners rapidly classified vowel sounds along an acoustic-phonetic continuum. We used support vector machine (SVM) classifiers and stability selection to determine when and where in the brain CP was best decoded across space and time via source-level analysis of the event related potentials (ERPs). We found that early (120 ms) whole-brain data decoded speech categories (i.e., prototypical vs. ambiguous speech tokens) with 95.16% accuracy [area under the curve (AUC) 95.14%; F1-score 95.00%]. Separate analyses on left hemisphere (LH) and right hemisphere (RH) responses showed that LH decoding was more robust and earlier than RH (89.03% vs. 86.45% accuracy; 140 ms vs. 200 ms). Stability (feature) selection identified 13 regions of interest (ROIs) out of 68 brain regions (including auditory cortex, supramarginal gyrus, and Brocas area) that showed categorical representation during stimulus encoding (0-260 ms). In contrast, 15 ROIs (including fronto-parietal regions, Broca’s area, motor cortex) were necessary to describe later decision stages (later 300 ms) of categorization but these areas were highly associated with the strength of listeners’ categorical hearing (i.e., slope of behavioral identification functions). Our data-driven multivariate models demonstrate that abstract categories emerge surprisingly early (∼120 ms) in the time course of speech processing and are dominated by engagement of a relatively compact fronto-temporal-parietal brain network.

## 1. INTRODUCTION

The human brain can map an incredibly large number of stimulus features into a smaller set of groups (Chang et al., 2010; Holt & Lotto, 2010), a process known as categorical perception (CP). Categories allow listeners to extract, manipulate, and precisely respond to sounds (C. T. Miller & Cohen, 2010; E. K. Miller et al., 2002, 2003; Russ et al., 2007; Tsunada & Cohen, 2014) despite wide variability in their acoustic properties. CP emerges in early life (Eimas et al., 1971) but is further modified by native language experience (Bidelman & Lee, 2015a; Kuhl et al., 1992; Xu et al., 2006). As such, CP plays an important role in understanding receptive communication and the building blocks of speech perception and language processing across the lifespan.

Event-related potentials (ERPs) are particularly useful for examining the brain mechanisms of phoneme and speech perception (Celsis et al., 1999; Molfese et al., 2005) given their excellent temporal resolution and the rapid time course required to process speech signals. Indeed, several ERP studies have documented neural correlates for CP via the ERPs (Bidelman, 2015; Binder et al., 2004; Chang et al., 2010). In particular, several studies have shown that the efficiency of listeners’ speech categorization varies in accordance with their underlying brain activity (Bidelman et al., 2013a; Bidelman & Alain, 2015; Bidelman & Lee, 2015b; Perlovsky, 2011). For example, Bidelman et al. demonstrated that brain responses in the time frame of 180- 320 ms were more robust for phonetic prototypes vs. ambiguous speech tokens, thereby reflecting category-level processing (Bidelman et al., 2020a). Other studies have shown links between N1-P2 amplitudes of the auditory cortical ERPs and the strength of listeners’ speech identification (Bidelman & Walker, 2017a) and labeling speeds (Al-Fahad et al., 2020) in speech categorization tasks (Bidelman et al., 2014; Bidelman & Alain, 2015). These findings are consistent with the notion that the early N1 and P2 waves of the ERPs are highly sensitive to speech processing and auditory object formation that is necessary to map sounds to meaning (Alain, 2007; Bidelman et al., 2013b; Wood et al., 1971).

The neural organization of speech categories also varies spatially, recruiting a widely distributed system across a number of brain regions. Neural responses are elicited by prototypical speech sounds (i.e., those heard with a strong phonetic category) differentially engage Heschl’s gyrus (HG) and inferior frontal gyrus (IFG) compared to ambiguous speech depending on a listeners perceptual skill level (Bidelman et al., 2013b; Bidelman & Lee, 2015a; Bidelman & Walker, 2017b; Mankel et al., 2020). This suggests emergent categorical representations within the early auditory-linguistic pathways. Similarly, Alho et al. found that category-specific representations were activated in left IFG (Alho et al., 2016) at an early-latency (115-140 ms). Collectively, in terms of the time course of processing, M/EEG studies agree that the neural underpinnings of speech categories emerge within the first few hundred milliseconds after stimulus onset and reflect abstract “category level-effects”(Toscano et al., 2018) and “phonemic categorization” (Liebenthal et al., 2010a).

Beyond conventional auditory-linguistic brain regions, neuroimaging also demonstrates a variety of additional areas important to speech perception and language processing (Hickok et al., 2011; Lee et al., 2012; Novick et al., 2010). Among them, superior partial lobe is associated with writing (Menon & Desmond, 2001) and supramarginal gyrus with phonological processing (Deschamps et al., 2014; Oberhuber et al., 2016) during speech and verbal working tasks. Relevant to CP, several studies have found that the left inferior parietal lobe is more activated during auditory phoneme sound categorization (Desai et al., 2008; Dufor et al., 2007; Husain et al., 2006). Indeed, auditory categorical processing has been shown to recruit superior temporal gyrus/sulcus, middle temporal gyrus, premotor cortex, inferior parietal cortex, planum temporal, and inferior frontal gyrus (Bidelman & Walker, 2019a; Guenther et al., 2004). Some other neuroimaging and electrocorticography studies have however shown that rostral anterior cingulate cortex is associated with speech control (Paus et al., 1993; Sahin et al., 2009; Tankus et al., 2012) and the orbitofrontal cortex in speech comprehension (Sabri et al., 2008). Under some circumstances (e.g., highly skilled listeners), speech categories can emerge as early as auditory cortex (Bidelman & Lee, 2015b; Bidelman & Walker, 2019a; Chang et al., 2010).

While category representations seem to emerge early in the time course of speech perception, the task of categorizing sounds can be further separated into pre- and post-perceptual stages of processing (i.e., stimulus encoding vs. decision mechanisms). “Early” vs. “late” stage models of category formation have long been discussed in the literature (Fox, 1984; McClelland & Elman, 1986; Noe & Fischer-Baum, 2020; Norris et al., 2000). However, few empirical studies have actually separately examined encoding and decision stages of CP. The human brain encodes speech stimuli within ∼250 ms after stimulus onset (Masmoudi et al., 2012) and decodes ∼300 ms after stimulus onset (Domenech & Dreher, 2010; Mostert et al., 2015). Previous studies have largely focused on these specific time windows (e.g., ERP waves) and brain regions when attempting to describe the neural basis of CP. While informative, such hypothesis-based testing can be restrictive and potentially miss the broader and distributed networks associated with speech-language processing that unfold on different time scales (Du et al., 2016; Rauschecker & Scott, 2009).

In this regard, machine learning (ML) techniques are increasingly used to “decode” high dimensional neuroimaging data and better understand different states of brain functionality as measured via EEG. ML is a branch of artificial intelligence that *“learns a model”* from the past data to predict future data (Cruz & Wishart, 2006). Moreover, data mining approaches in ML identify important properties in neural activity with high accuracy without intervention from human observers. It would be meaningful if brain functioning that has been linked with speech processing (e.g., CP) could be decoded from neural data without, or at least with minimal, *a priori* assumptions on when and where those representation emerge. Indeed, laying the groundwork for the present work, we have recently shown that the speed of listeners’ identification in speech categorization tasks can be directly decoded from their full-brain EEGs using an entirely data-drive approach (Al-Fahad et al., 2020). We have also shown that ML can decode age-related changes in speech processing that occur in older adults (Mahmud et al., 2020).

Departing from previous hypothesis-driven studies (Bidelman & Alain, 2015; Bidelman & Walker, 2019a, 2017a), the current work used a comprehensive, data-driven approach to examine the neural mechanisms of speech categorization during encoding and decision stages of processing using whole-brain, electrophysiological data. We analyzed speech-evoked ERPs from high density EEGs recorded during a rapid speech categorization task in young, normal hearing listeners. Our approach applied state-of-the-art ML techniques including neural classifiers and feature selection methods (i.e., stability selection) to source-level ERPs to investigate the spatiotemporal dynamics of speech categorization. We aimed to determine when and where neural activity from full-brain EEGs differentiated phonetic from phonetically ambiguous speech sounds, and thus showed the strongest evidence of categorical processing using an entirely data-driven, machine learning approach.

## 2. MATERIALS & METHODS

### 2.1 Participants

Forty-nine young adults (male: 15, female: 34; aged 18 to 33 years) were recruited as participants from the University of Memphis student body to participate into our ongoing studies on the neural basis of speech perception and auditory categorization (Bidelman et al., 2020b; Bidelman & Walker, 2017b; Mankel et al., 2020). All participants had normal hearing sensitivity (i.e., *<*25 dB HL between 500-2000 Hz). Listeners were-right handed (Oldfield, 1971) and had achieved a collegiate level of education. None reported any history of neurological disease. All participants were paid for their time and gave informed written consent in accordance with the declaration of Helsinki and a protocol approved by the Institutional Review Board at the University of Memphis.

### 2.2 Stimuli & task

We used a synthetic five-step vowel token continuum to assess the most discriminating spatiotemporal features while categorizing prototypical vowel speech from ambiguous speech (Bidelman et al., 2013b, 2014). Speech spectrograms are represented in Fig. 1A. Each token of the continuum was separated by equidistant steps acoustically based on the first formant frequency (F1) and perceived to categorically from /u/ to /a/. Each speech token was 100 ms, including 10 ms rise/fall to minimize the spectral splatter in the stimuli. Each speech token contained an identical voice fundamental frequency (F0), second (F2), and third formant (F3) frequencies (F0:150 Hz, F2: 1090 Hz, and F3:2350 Hz). To create a phonetic continuum that varied in percept from /u/ to /a/, F1 frequency was parameterized over five equal steps from 430 Hz to 730 Hz (Bidelman et al., 2013b).

**Figure 1:**
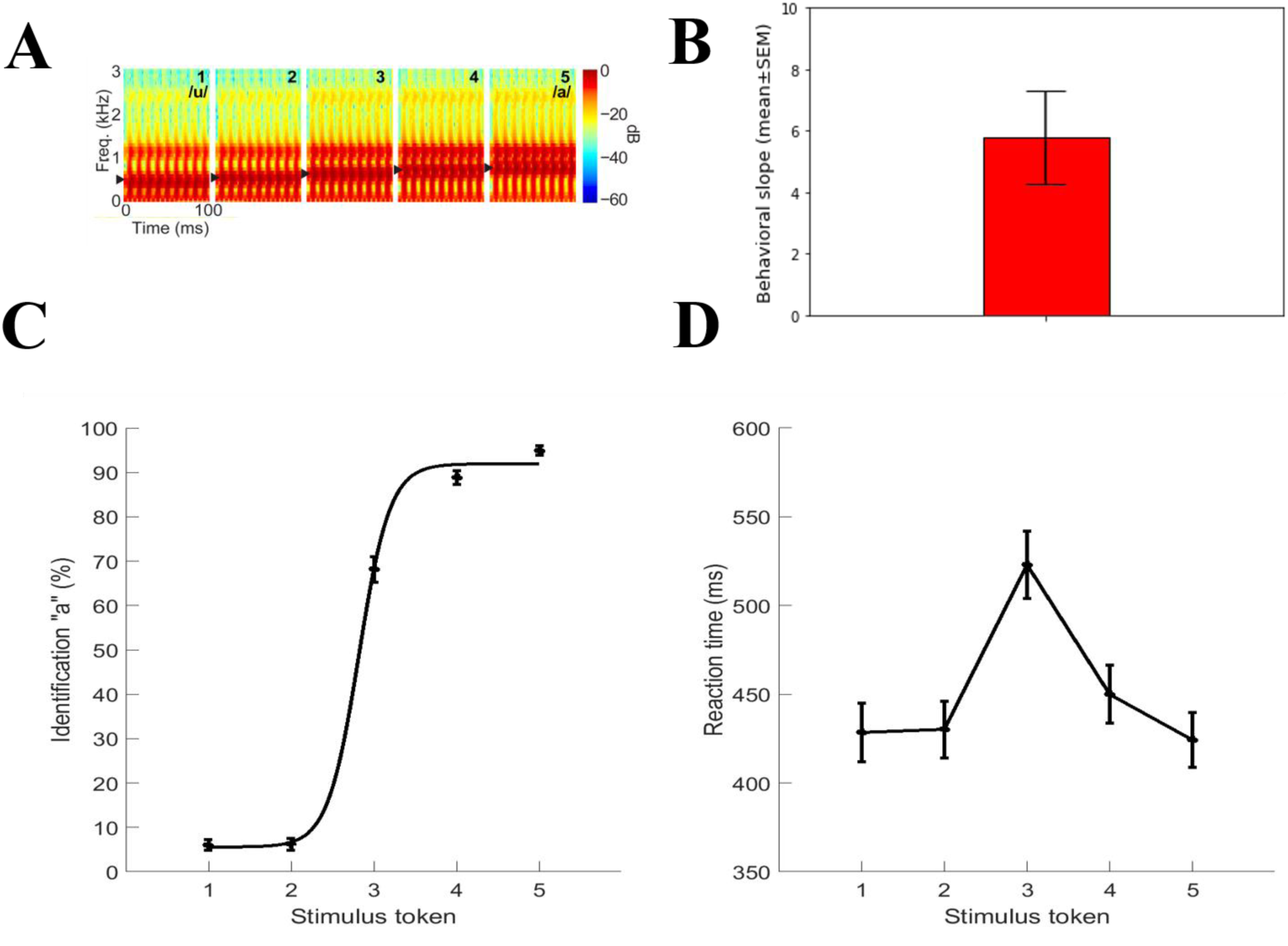
Speech stimuli and behavioral results. **A)** Acoustic spectrograms of the speech continuum from */u/* and */a/*. **B)** Behavioral slope. **C)** Psychometric functions showing % “a” identification of each token. Listeners’ perception abruptly shifts near the continuum midpoint, reflecting a flip in perceived phonetic category (i.e., “u” to “a”). **D)** Reaction time (RT) for identifying each token. RTs are fastest for category prototypes (i.e., Tk1/5) and slow when classifying ambiguous tokens at the continuum midpoint (i.e., Tk3). Errorbars = ±1 s.e.m.

Stimuli were presented binaurally at an intensity of 83 dB SPL through earphones (ER 2; Etymotic Research). Participants heard each token 150-200 times presented in random order. They were asked to label each sound in a binary identification task (“/u/” or “/a/”) as fast and accurately as possible. Their response and reaction time were logged. The interstimulus interval (ISI) was jittered randomly between 400 and 600 ms (20 ms step and rectangular distribution) following listeners’ behavioral responses to avoid anticipating the next trial (Luck, 2005).

### 2.3 EEG recordings and data pre-procedures

During the behavioral task, EEG was recorded from 64 channels at standard 10-10 electrode locations on the scalp (Oostenveld and Praamstra 2001). Continuous EEGs were digitized using Neuroscan SynAmps RT amplifiers at a sampling rate of 500 Hz. Subsequent preprocessing was conducted in the Curry 7 neuroimaging software suite, and customized routines coded in MATLAB. Ocular artifacts (e.g., eye-blinks) were corrected in the continuous EEG using principal component analysis (PCA) (Picton et al., 2000) and then filtered (1-100 Hz bandpass; notched filtered 60 Hz). Cleaned EEGs were then epoched into single trials (−200 to 800 ms, where *t* = 0 was stimulus onset).

### 2.4 EEG source localization

To disentangle the sources of CP-related EEG activity, we reconstructed the scalp-recorded responses by performing a distributed source analysis in the Brainstorm software package (Tadel et al., 2011). All analyses were performed on single-trial data. We used a realistic boundary element head model (BEM) volume conductor and standard low-resolution brain electromagnetic tomography (sLORETA) as the inverse solution within Brainstorm (Tadel et al., 2011). A BEM model has less spatial errors than other existing head models (e.g., concentric spherical head model). We used Brainstorm’s default parameter settings (SNR=3.00, regularization noise covariance = 0.1). From each single-trial sLORETA volume, we extracted the time-courses within 68 functional regions of interest (ROIs) across the left and right hemispheres defined by the Desikan-Killiany (DK) atlas (Desikan et al., 2006) (LH: 34 ROIs and RH: 34 ROIs). Single-trial data were then baseline corrected to the epoch’s pre-stimulus interval (−200-0 ms).

Since we were interested to decode prototypical (Tk1/5) from ambiguous speech (Tk3)— a marker of categorical processing (Bidelman, 2015; Bidelman & Walker, 2019b; Liebenthal et al., 2010b)—we merged Tk1 and Tk5 responses since they reflect prototypical vowel categories (“u” vs. “a’). In contrast, Tk3 reflects a bistable percept—an category-ambiguous sound listeners sometimes label as “u” or “a” (Bidelman et al., 2020a; Bidelman & Walker, 2017b; Mankel et al., 2020). To ensure an equal number of trials and signal to noise ratio (SNR) for prototypical and ambiguous stimuli, we considered only 50% of the data from the merged (Tk1/5) samples.

### 2.5 Feature extraction

Previous computational studies have found that ERPs averaged over 100 trials provided the best classification of data while maintaining reasonable signal SNR and computational efficiency (Al-Fahad et al., 2020; Mahmud et al., 2020). We quantified source-level ERPs with a mean bootstrapping approach (James et al., 2013) by randomly averaging over 100 trials (with replacement) 30 times (Al-Fahad et al., 2020) for each stimulus condition per participant. For each resample and ROI time course, we measured the mean amplitude within a 20 ms sliding window (without overlapping) in the post-stimulus interval (i.e., 0 to 800 ms). In post hoc analysis, we parsed the epoch into “encoding” (0-260 ms) and “decoding/decision process” intervals (>300 ms) to investigate neural decoding related to pre- and post-perceptual processing, respectively. The sliding window resulted in 40 (800ms/20ms) ERP features (i.e., mean amplitude per window) for each ROI waveform, yielding a total of 68*40=2720 features per token (e.g., Tk1/5 vs. Tk3) from each listeners’ data. Thus, the encoding and decision windows contained 13*68=884 (encoding) and 25*68=1700 (decision) ERP features. ERPs features were then used as input to an SVM classifier to access the temporal dynamics of the data and determine when in time CP was decodable from brain activity. State-of-the art variable selection (stability selection; see *Section 2.7*) (Meinshausen & Bühlmann, 2010) was then applied for identifying where in the brain (e.g., which ROIs) were involved in encoding and decision processes with regard to the categorization of speech. Before submitting to the SVM classifier, the data were z-score normalized to ensure all features were on a common scale range (Casale et al., 2008).

### 2.6 SVM classification to identify temporal dynamics of CP

Parameter optimized Support Vector Machine (SVM) classifiers provide better performance with small sample sizes data which is common in human neuroimaging studies. Classifier performance is greatly affected by tunable parameters in the SVM model (e.g., kernel, C, *γ*)^1^ (Hsu et al., 2003). To avoid bias in parameter selection, we used a grid search approach during the training phase to find optimal kernel, *C*, and *γ* values. We randomly split the data into training (80%) and test (20%) sets (Park et al., 2011). During the training phase (e.g., using 80% data), we fine-tuned the *C*, and *γ* parameters using grid search to find the optimal values such that the resulting classifier accurately distinguished prototypical vs. ambiguous speech in the test data that models never seen. The grid search process was conducted with five-fold cross validation, kernels = ‘RBF’, fine-tune 20 different values of (*C* and *γ*) in the following range *C* = [1e-2 to 1e3], and *γ* = [1e-4 to 1e2] (Mahmud et al., 2020). The SVM learned the support vectors from the training data that comprised the attributes (e.g., ERP features) and class labels (e.g., Tk1/5 vs. Tk3). Then we selected the best model that has maximum margin with the optimal value of *C* and *γ* for predicting the unseen test data (only by providing the attributes but no class labels). The classification performance metrics (accuracy, F1-score, precision, and recall) are calculated from standard formulas (Saito & Rehmsmeier, 2015).

### 2.7 Stability selection to identify spatial dynamics of CP

Our data included a large number (∼2700) of ERP measurements for each stimulus condition of interest (e.g., Tk1/5 vs. Tk3). Larger numbers of variable/features can lead to overfitting and weak generalization in classification problems since the majority of features from brain activity (i.e., different ROIs, time segments) do not provide discriminative power for decoding CP. Consequently, we aimed to select a limited set of the most salient discriminating features. Stability selection is a state-of-the art feature selection method that works well in high dimensional or sparse data based on the Lasso (least absolute shrinkage and selection operator) (Meinshausen & Bühlmann, 2010; Yin et al., 2017). Stability selection can identify the most stable (relevant) features out of a large number of features over a range of model parameters, even if the necessary conditions required for the original Lasso method are violated (Meinshausen & Bühlmann, 2010).

In stability selection, a feature is considered to be more stable if it is more frequently selected over repeated subsampling of the data (Nogueira et al., 2017). Basically, the Randomized Lasso randomly subsamples the training data and fits a L1 penalized logistic regression model to optimize the error. Over many iterations, feature scores are (re)calculated. The features are shrunk to zero by multiplying the features’ co-efficient by zero while the stability score is lower. Surviving non-zero features are considered important variables for classification. Detailed interpretation and mathematical equations of stability selection are explained in (Meinshausen & Bühlmann, 2010). The stability selection solution is less affected by the choice of the initial regularization parameters. Consequently, it is extremely general and widely used in high dimensional data even when the noise level is unknown.

In our implementation of stability selection, we used a sample fraction = 0.75, number of resamples = 1000, and tolerance = 0.01 (Meinshausen & Bühlmann, 2010). In the Lasso algorithm, the feature scores were scaled between 0 to 1, where 0 is the lowest score (i.e., irrelevant feature) and 1 is the highest score (i.e., most salient or stable feature). We estimated the regularization parameter from the data using the least angle regression (LARs) algorithm (Efron et al., 2004; Friedman et al., 2010). Over 1000 iterations, Randomized Lasso provided the overall feature scores (0∼1) based on the number of times a variable was selected. We ranked stability scores to identify the most important, consistent, stable, and invariant features that could decode speech categories via the EEG (i.e., correctly classify Tk1/5 vs. Tk3). We submitted these ranked features and corresponding class labels to an SVM classifier with different stability thresholds and observed the model performance.

## 3. RESULTS

### 3.1 Behavioral results

Behavioral identification (%) functions and reaction time (ms) for speech categorization are depicted in Fig. 1C, Fig. 1D, respectively. Listeners responses abruptly shifted in speech identity (/u/ vs. /a/) near the midpoint of the continuum, reflecting a change in perceived category. The behavioral speed of speech labeling (e.g., reaction time (RT)) were computed listeners’ median response latency for a given condition across the all trials. RTs outside of 250-2500 ms were deemed outliers and excluded from further analysis (Bidelman et al., 2013a; Bidelman & Walker, 2017a). Listeners spent more time classifying the ambiguous (Tk3) than prototypical speech tokens (e.g., Tk1/5), further confirming categorical hearing (Pisoni & Tash, 1974). For each continuum, the identification scores were fit with a two parameters sigmoid function; 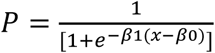, where *P* is the proportion of the trial identification as a function of a given vowel, *x* is the step number along the stimulus continuum, and *β0* and *β1* the location and slope of the logistic fit estimated using the nonlinear least-squares regression (Bidelman et al., 2014; Bidelman & Walker, 2017a). The slopes of listeners’ sigmoidal psychometric function, reflecting the strength of their CP, is presented in Figure 1B.

### 3.2 Decoding the time-course of speech categorization from ERPs

We first examined how well categorical speech information could be decoded from whole-brain and individual hemisphere (e.g., LH and RH) ERPs data. During pilot modeling, we carried the grid search approach (mentioned in method). The optimal values of C and γ parameters corresponding to the maximum speech decoding reported in Table 1 were: [C=10, γ=0.05 for whole-brain data; C=20, γ=0.01 for LH data; C=20, γ=0.01 for RH data]. We then selected the best model and predicted the class labels (e.g., Tk1/5 vs. Tk3) by feeding the feature vectors only from the unseen test data. The performance metrics were calculated from predicted class labels and true class labels. Time-varying accuracy of the SVM classifier (i.e., distinguishing Tk1/5 vs. Tk3 responses) is shown in Figure 3.

**Table 1:** Performance metrics of the SVM classifier corresponding to maximal decoding of prototypical vs. ambiguous vowels from ERPs.

| Metric (%) | Whole-brain features | LH features | RH features |
| --- | --- | --- | --- |
| Accuracy | 95.16 | 89.03 | 86.45 |
| AUC | 95.14 | 89.18 | 86.45 |
| F1-score | 95.00 | 89.00 | 86.00 |
| Precision | 95.00 | 89.00 | 87.00 |
| Recall | 95.00 | 89.00 | 86.00 |

**Figure 2:**
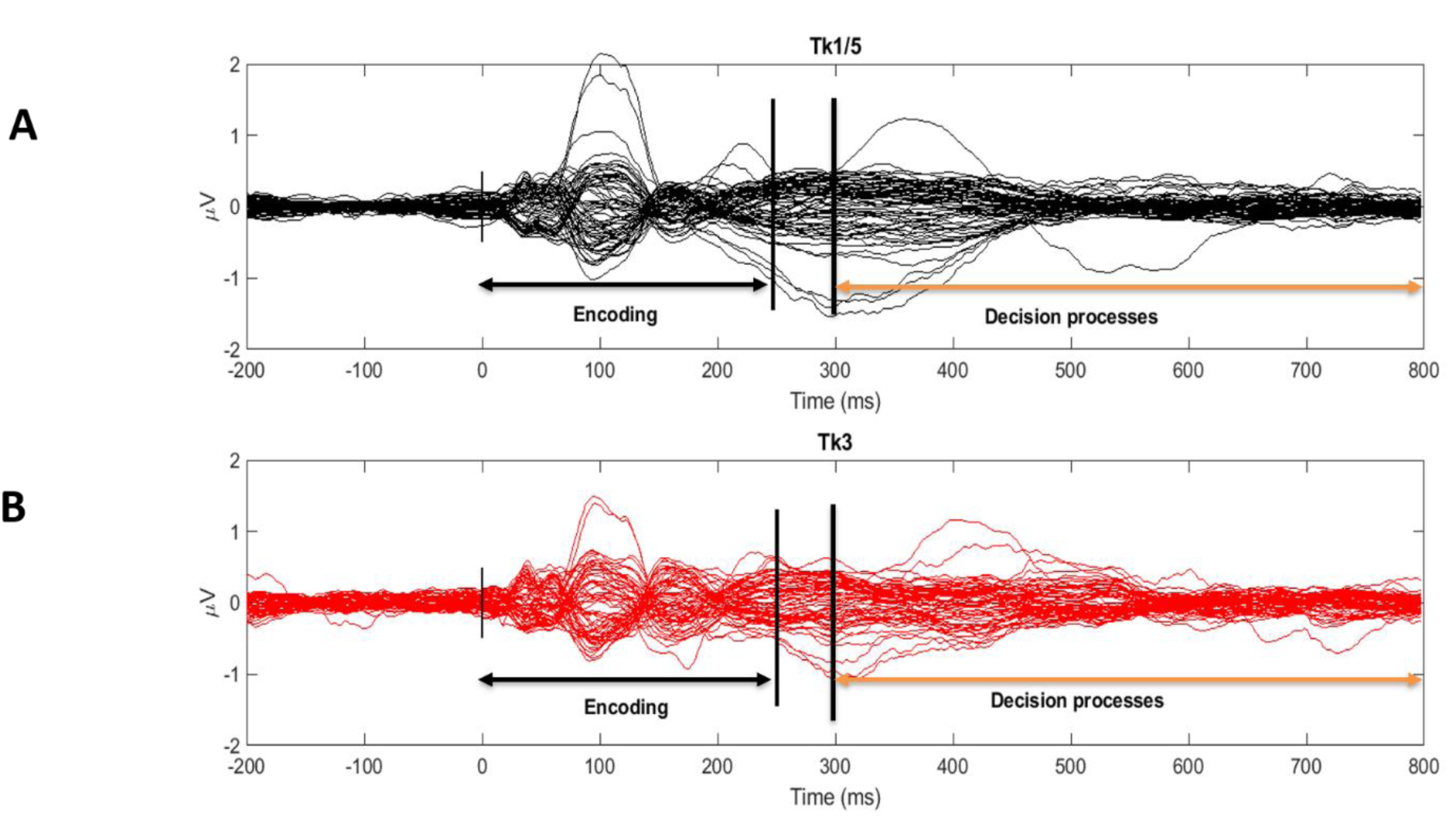
Grand averaged butterfly plots of scalp ERPs (64 channels) to prototypical (**A**; Tk1/5) vs. category-ambiguous (**B**; Tk3) vowels. Vertical lines demarcate segments for the stimulus encoding (0-260 ms) and decision period (300 ms-800 ms) analysis windows. *t*=0 marks stimulus onset.

**Figure 3:**
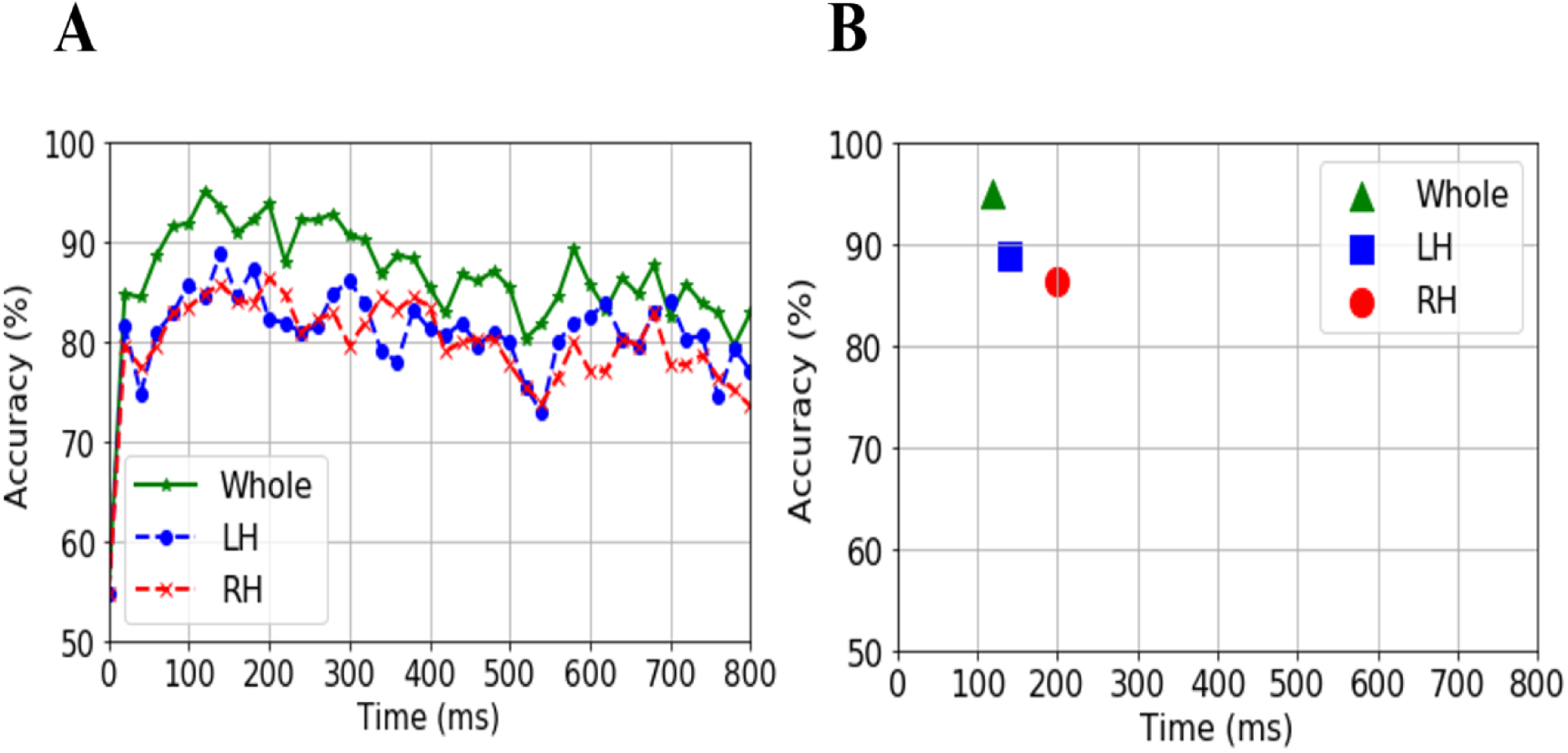
SVM classifier accuracy decoding speech categories from source ERPs. **A)** Decoding using whole-brain vs. hemispheres-specific data (LH and RH) across the epoch window. **B)** Maximum classifier accuracy was observed at ∼120 ms suggesting category representations emerge early, ∼200 ms before listeners’ behavioral categorization decisions (cf. Fig. 1C).

Decoding was generally at chance level (54%) at stimulus onset (i.e., t = 0) but increased rapidly to a maximum accuracy of 95.16% by 120 ms. The individual hemispheres alone were less accurate and decoded speech categories later in time compared to whole-brain data (LH: 89.03% at 140 ms; RH: 86.45% at 200 ms) (Fig. 3B). Other important performance metrics of the SVMs at maximum decoding are reported in Table 1. Collectively, the earlier and improved ability of LH compared to RH in decoding phonetic categories is consistent with a left hemisphere bias in speech and language processing (Hickok & Poeppel, 2000). More critically, the early time course of decoding (120-150 ms) confirms that category level information, an abstract code, emerges very early in the neural chronometry of speech processing and well before listeners’ execute their behavioral decision (cf. reaction times in Fig. 1D) (Alho et al., 2016; Bidelman et al., 2013c; de Taillez et al., 2020).

### 3.3 Decoding the spatial regions underlying categorization: stimulus encoding vs. decision

We used stability selection to find the most critical brain ROIs that were associated with categorical organization in the encoding (pre-perceptual) vs. decision (post-perceptual) periods of the task structure (see Fig. 2). ERP features were considered stable (relevant) if they yielded a decoding accuracy performance >80%. The effect of stability threshold selection in the encoding and decision windows is illustrated in Figure 4. Each bin of histogram demonstrates the number of features in a range of stability threshold. The x-axis has four labels. The first line represents the stability score (0 to 1); the second and third line show the number of features and percentage of the selected features in the corresponding bin; line four represents the cumulative unique ROIs up to the lower boundary of the bin. The solid black and dotted red semi bell-shaped curves of Figure 4 represent classification accuracy and AUC, respectively for the different stability thresholds. In this analysis, the number of unique brain ROIs represents distinct functional brain ROIs of the DK atlas and the number of features represents different time windows extracted from source ERPs. Features selected at each stability threshold were then submitted to an SVM classifier separately for the stimulus encoding and response decision periods.

**Figure 4:**
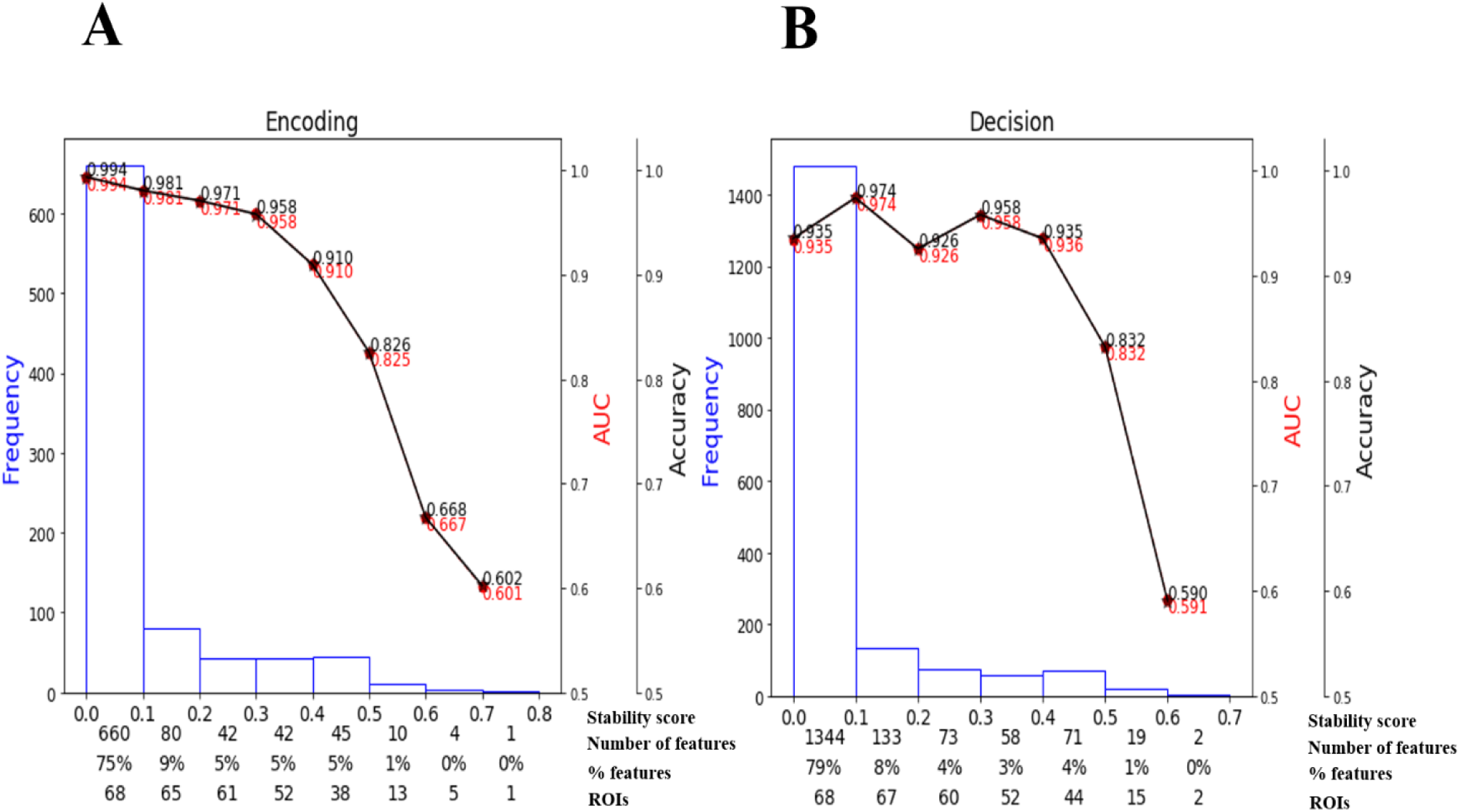
Effect of stability score threshold on model performance during (**A**) encoding and (**B**) decision period of the CP task. The bottom of the x-axis has four labels; *Stability score* represents the stability score range of each bin (scores: 0∼1); *Number of features*, number of features under each bin; *% features*, the corresponding percentage of selected features; *ROIs*, number of cumulative unique brain regions up to the lower boundary of the bin.

During stimulus encoding (0-260 ms), 75% of features yielded stability scores 0 to 0.1. Thus, the majority of spatiotemporal ERP features were selected less than 10% out of 1000 model iterations and therefore carry weak importance in terms of describing categorical speech processing during stimulus encoding. In contrast, at a more conservative stability score of 0.3, 102 (11%) out of 884 ERP features selected from 52 ROIs were able to encode prototypical from ambiguous speech at near-ceiling accuracy (95.8%). Accuracy decreased precipitously with higher (more conservative) stability thresholds resulting in fewer (though more informative) brain ROIs describing category processing. For example, a stability score of 0.6—selecting only the most behaviorally-relevant features—still encoded speech categories well above chance (66.8%) with only 5 features from 5 ROIs. At stability score 0.5, speech encoding accuracy 82.6% only using 15 features from 13 unique ROIs. A BrainO visualization (Moinuddin et al., 2019) of relevant ROIs for the encoding period (threshold stability score ≥ 0.5) is shown in Figure 5 with additional details in Table 2.

**Table 2:** Most important brain regions describing speech categorization during stimulus encoding (13 ROIs) and response decision (15 ROIs) at a stability threshold ≥ 0.5.

| Rank | Encoding (82.6% total accuracy) |  |  | Decision (83.2% total accuracy) |  |  |
| --- | --- | --- | --- | --- | --- | --- |
|  | ROI name | ROI abbrev. | Stability score | ROI name | ROI abbrev. | Stability score |
| 1 | Supramarginal L | ISUPRA | 0.73 | Superior parietal L | ISP | 0.63 |
| 2 | Caudal anterior cingulate R | rCAC | 0.66 | Insula L | IINS | 0.60 |
| 3 | Inferior parietal L | IIP | 0.65 | Isthmus cingulate R | rIST | 0.58 |
| 4 | Pars orbitalis R | rPOB | 0.61 | Pars opercularis R | rPOP | 0.58 |
| 5 | Transverse temporal L | ITRANS | 0.61 | Superior frontal L | ISF | 0.57 |
| 6 | Superior frontal R | rSF | 0.58 | Caudal middle frontal R | rCMF | 0.57 |
| 7 | Pars opercularis L | IPOP | 0.57 | Isthmus cingulate L | IIST | 0.56 |
| 8 | Lateral orbitofrontal L | ILOF | 0.57 | Pars triangularis R | rPT | 0.54 |
| 9 | Superior frontal L | ISF | 0.55 | Caudal middle frontal L | ICMF | 0.54 |
| 10 | Pars triangularis R | rPT | 0.54 | entorhinal L | IENT | 0.53 |
| 11 | Superior parietal R | rSP | 0.53 | Pars opercularis L | IPOP | 0.53 |
| 12 | Caudal middle frontal R | rCMF | 0.53 | Paracentral R | rPARAC | 0.52 |
| 13 | fusiform L | IFUS | 0.52 | Inferior parietal L | IIP | 0.51 |
| 14 |  |  |  | Parahippocampal R | rPHIP | 0.51 |
| 15 |  |  |  | Postcentral L | IPOC | 0.51 |

**Figure 5:**
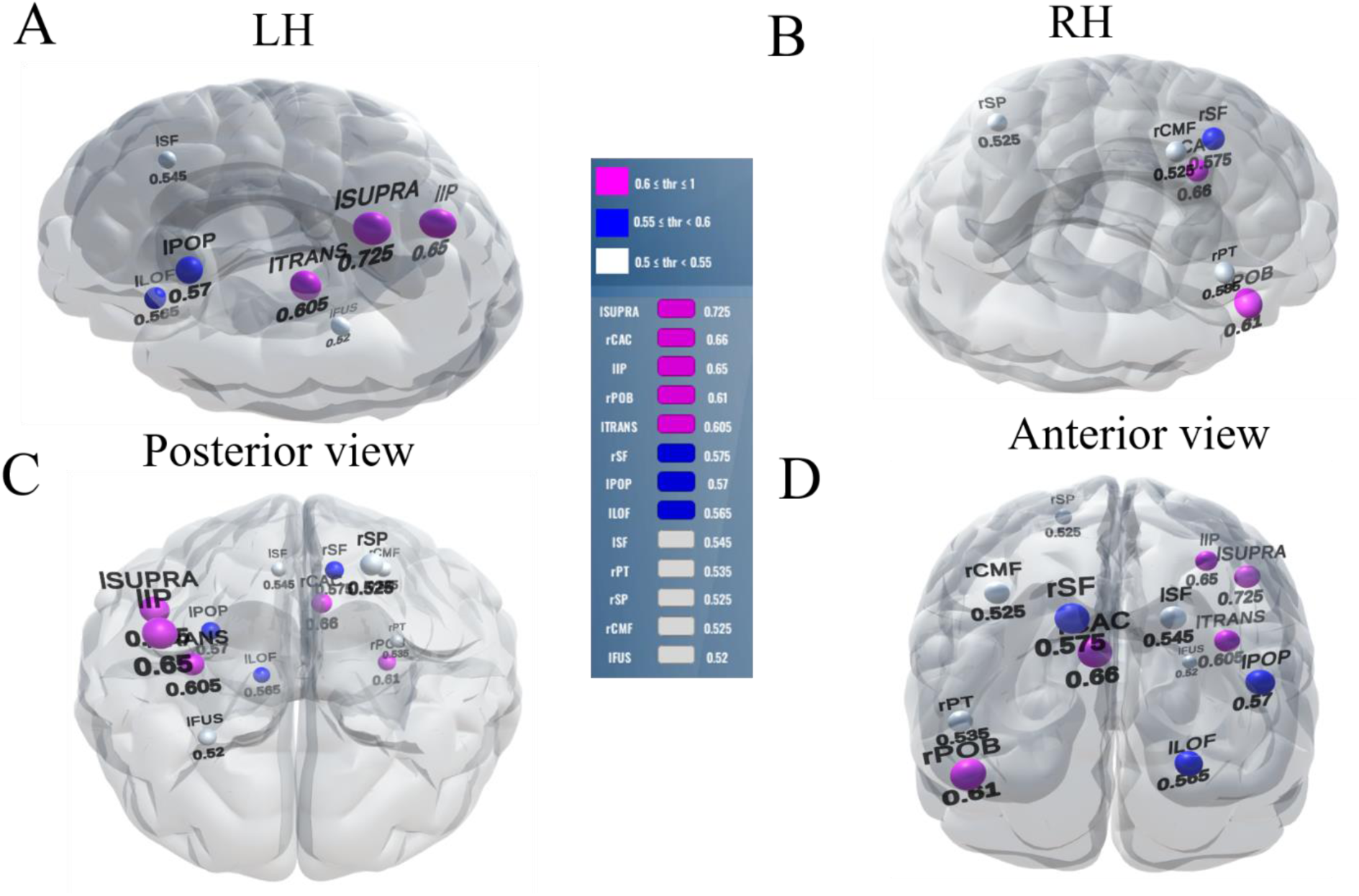
Stable (most consistent) neural network during the *encoding period* of CP. Visualization of brain ROIs corresponding to ≥ 0.50 stability threshold (13 top selected ROIs which show categorical organization (e.g., Tk1/5 ≠ Tk3) at 82.6%. **(A)** LH **(B)** RH **(C)** Posterior view (**D)** Anterior view. Color legend demarcations show high (pink), moderate (blue), and low (white) stability scores. l/r = left/right; SUPRA, supramarginal; CAC, caudal anterior cingulate; IP, inferior parietal; POB, pars orbitalis; TRANS, transverse temporal; SF, superior frontal; POP, pars opercularis; LOF, lateral orbitofrontal; PT, pars triangularis; SP, superior parietal; CMF, caudal middle frontal; FUS, fusiform.

**Figure 6:**
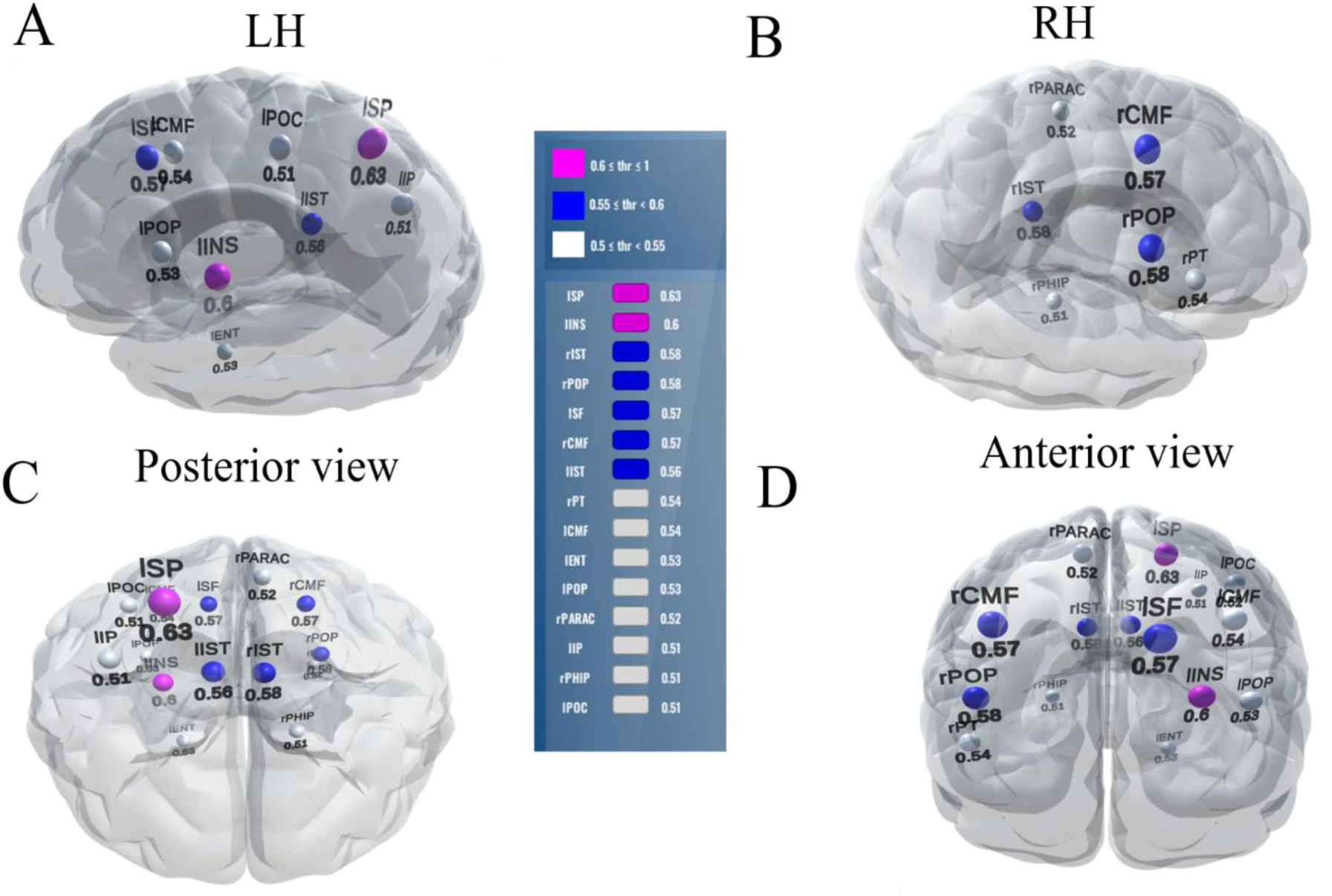
Stable (most consistent) neural network during the *decision period* of CP. Visualization of brain ROIs corresponding to ≥ 0.50 stability threshold (15 top selected ROIs which decode Tk1/5 from Tk3 at 83.2%. Otherwise as in Figure 5. SP, superior parietal; INS, Insula; POP, pars opercularis ; SF, superior frontal; CMF, caudal middle frontal; IST, isthmus cingulate; PT, pars triangularis; CMF, caudal middle frontal; ENT, entorhinal; PARAC, paracentral; IP, inferior parietal; PHIP, para hippocampal ;POC, postcentral.

During the decision period following stimulus encoding (> 300 ms), corresponding to the stability score 0.4, only 92 (5%) out of 1700 ERP features were selected, and the classifier showed decoding accuracy of 93.5% (AUC 93.6%). At a stability score 0.5 (corresponding to 83.2% accuracy), only 21 (1%) out 1700 ERP features from 15 unique ROIs were needed to describe categorical processing.

### 3.4 Brain-behavior correspondences

Multivariate regression analysis is widely used to investigate when more than one predictor simultaneously influences an outcome variable (Hanley, 1983; Royston & Sauerbrei, 2008). To evaluate the behavioral relevance of the brain regions identified via stability selection, we conducted multivariate regression using weighted least squares (WLS) regression (Ruppert & Wand, 1994). Regressions were computed between the 15 ROI ERPs identified in the decision interval and listeners’ behavioral slopes (Fig. 1B), which indexes their degree of categorical hearing. We computed the mean neural response (i.e., ERP) within each selected region across the stimuli [mean ERP of (Tk1/5 & Tk3)] and then regressed the 15 ROI responses simultaneously against listeners’ behavioral slope. The inverse of the absolute error values of the ordinary least squares were used as weights in the WLS to reduce the effect of heteroscedasticity (Seabold & Perktold, 2010; *Weighted Regression in SAS, R, and Python*, n.d.). The multivariate model robustly predicted listeners’ behavioral CP from neural data (R^2^ = 0.85, *p*<0.00001; Table 3), demonstrating the selected 15 ROIs identified via ML (i.e., stability selection) carried behaviorally relevant information regarding CP.

**Table 3:**
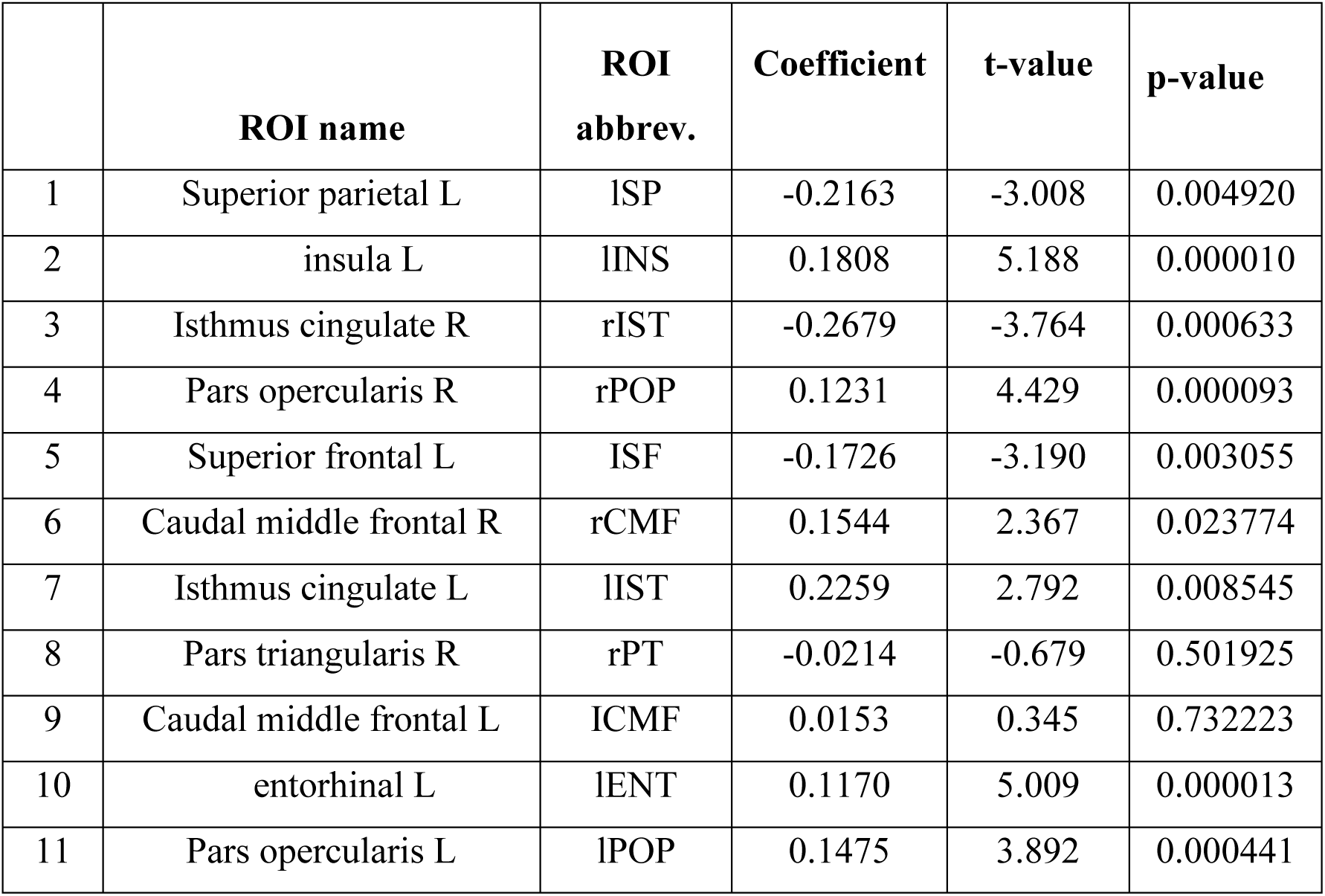
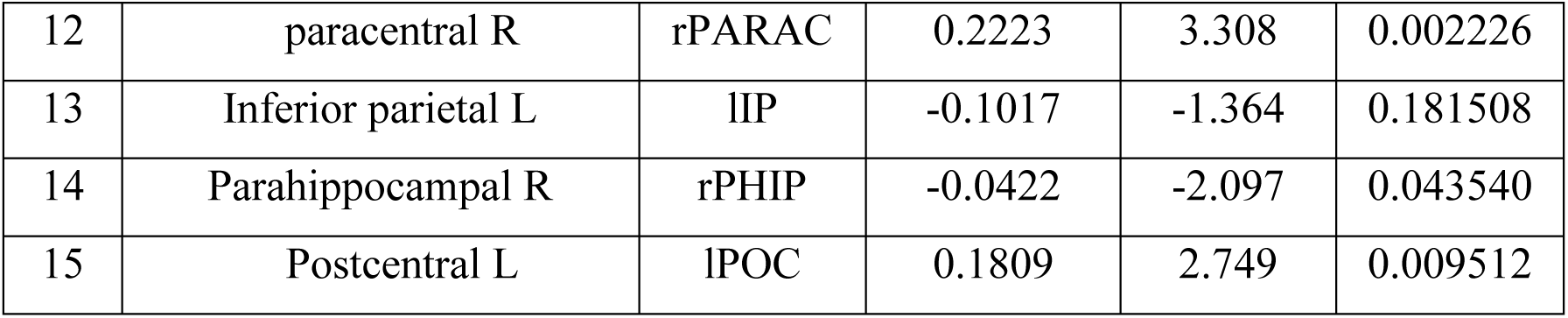
WLS regression results describing how individual brain ROIs predict behavioral CP.

## 4. DISCUSSION

We conducted machine learning analyses on EEG to examine the spatiotemporal dynamics of speech processing during rapid speech sound categorization. We found that speech categories are best decoded via patterned neural activity occurring within 120 ms and no later than 200 ms. We also identified the most relevant brain regions that are involved in encoding and decision stages of the categorization process. Our findings show a small set of brain areas (15 ROIs) robustly predicts listeners’ categorical decisions, accounting for 85.0% of the variance in behavior.

### 4.1 Speech categories are decoded early (<150 ms) in the time course of perception

We replicate and extend previous work by using whole-brain EEG and SVM neural classifiers to examine the time-course and hemispheric asymmetry as the brain decodes the identity of speech sounds. We found optimal speech decoding in the time frame of the N1 wave (120 ms) of the auditory ERPs using full-brain data. Analysis by hemisphere further showed that LH yielded better and earlier decoding than the RH, where optimal decoding occurred 20-80 ms later (LH: 140 ms; RH: 200 ms). These latencies are compatible with the N1-P2 waves of the auditory ERPs and suggest a rapid speed to phonetic categorization (Alho et al., 2016; Bidelman et al., 2013c; de Taillez et al., 2020). Our results are consistent with previous neuroimaging studies that have shown the N1 and P2 ERPs are sensitive to auditory perceptual object identification (Alain, 2007; Bidelman et al., 2013b; Wood et al., 1971). The better decoding by LH as compared to RH activity is consistent with the dominance of LH in phoneme discrimination and speech sound processing (Bidelman & Howell, 2016; Bidelman & Walker, 2019b; Frost et al., 1999; Tervaniemi & Hugdahl, 2003; Zatorre et al., 1992). Our neural decoding results also corroborate previous hypothesis-driven work (Bidelman et al., 2013c, 2014; Chang et al., 2010) by confirming speech sounds are converted to an abstract, categorical representation within the first few hundred milliseconds after stimulus onset

### 4.2 Differential brain-networks involved in encoding and decision processing

Our results help identify the most stable, relevant, and invariant functional brain ROIs that support the brain-networks involved in encoding and decision processes of speech categorization using an entirely data-driven approach (stability selection coupled with SVM). During stimulus encoding, stability selection have identified 13 consistent ROIs that differentiate speech categories (82.6% accuracy; 0.5 stability threshold). Out of these 13 regions, eight of the ROIs are critically involved in the dorsal-ventral pathway for speech-language processing (Hickok & Poeppel, 2004). These included areas in frontal lobe including Broca’s area [BA 44, (i.e., pars opercularis L, pars triangularis R)], three regions from parietal and two regions from temporal lobe including primary auditory cortex (i.e., transverse temporal L). For later decision stages of the task, the same criterion of decoding performance (83.2% @ 0.5 stability threshold) have identified 15 ROIs that showed categorical neural organization. Out of these 15 regions, eight areas are from frontal lobe including Broca’s area [BA 44, (i.e., pars opercularis L, pars opercularis R), and BA 45 (i.e., pars triangularis R)], four regions from parietal lobe, and three regions from temporal lobe. Our data reveal two, relatively sparse, and partially overlapping neural networks that support different stages of speech categorization process.

Among the encoding and decision networks identified from our EEG data, five regions were common between the two topologies. Notably were the inclusion of BA44/45 (i.e., canonical Brocas’ area) that are heavily involved in speech-language processing (Hickok et al., 2011; Lee et al., 2012; Novick et al., 2010). The left inferior parietal lobe also appears as a common hub among the two networks. Superior parietal areas have been linked with auditory, phoneme, sound categorization, particularly when listeners are asked to resolve context or ambiguity (Dufor et al., 2007; Feng et al., 2018; Myers & Blumstein, 2008). Involvement of superior frontal lobe in both networks is perhaps consistent with its role in higher cognitive functions and working memory (Klingberg et al., 2002; Nyberg et al., 2003). The fact that these extra-sensory regions can decode category structure even during stimulus encoding (< 150 ms) suggests that the formation of speech categories might operate nearly in parallel within lower-order (sensory) and higher-order (cognitive-control) brain structures (Toscano et al., 2018). However, these category representations need not be isomorphic across the brain. For example, category formation might reflect a cascade of events where speech units are reinforced and further discretized by a recontact of acoustic-phonetic with lexical representation of the speech category (Myers & Blumstein, 2008).

Notable among the non-overlapping regions between stages were left primary auditory cortex (transverse temporal) and supramarginal gyrus, both of which were exclusive to the stimulus encoding period. Both regions have been implicated in the early acoustic analysis of the speech signal and related phonological processing (Deschamps et al., 2014; Geiser et al., 2008; Hickok et al., 2000; Oberhuber et al., 2016; Whitwell et al., 2013; Zatorre et al., 1992). Intuitively, their absence during the decision stage further suggests the categorical representation of speech, while present early in time (< 150 ms), might take different forms in auditory-sensory cortex before being broadcast to decision mechanisms downstream.

Left postcentral gyrus is also exclusive during decision. Activation of this area proximal to the behavioral response execution most probably reflects motor planning and/or speech reconstruction (Martin et al., 2014). Additional non-overlapping ROIs included pars opercularis in the RH. Right IFG has been implicated in attentional control and response imbibition (Hampshire et al., 2010), which is consistent with its exclusive involvement in the decision stage of our task. Presumably, the other non-overlapping regions identified via stability selection (superior parietal L, insula L, Isthmus cingulate (l/rIST), caudal middle frontal L, entorhinal L, paracentral R, parahippocampal R) are also involved in decision processes, though as of yet, in an unknown way. Minimally, the involvement parahippocampal regions implies putative memory and retrieval processes. Still, more detailed localization studies (e.g., using fMRI) are needed to validate our EEG data, which offers a much coarser spatial resolution.

It is noticeable that during encoding, 7 out of 13 ROIs are from LH; for decoding, 9 out of 15 ROIs. The left hemisphere bias in our decoding data is perhaps expected given the LH dominance in auditory language processing (Caplan, 1994; Hull & Vaid, 2006; Tzourio et al., 1998). Moreover, our results support previous studies by confirming a bilateral fronto-parietal network involved in auditory attentional, working memory (Belin et al., 2002; Schneiders et al., 2012), sound discrimination tasks (Hickok & Poeppel, 2000), and phoneme categorization (Bidelman & Walker, 2019a; Lee et al., 2012; Loui, 2015). Interestingly, our study shows that only 15 brain regions (during decision) are needed to predict listeners’ behavior CP with 85.0% accuracy.

## 5. ACKNOWLEDGMENTS

Requests for data and materials should be directed to G.M.B. This work was supported by the National Institutes of Health (NIH/NIDCD R01DC016267) and department of Electrical and Computer Engineering at the University of Memphis.

## Footnotes

1 Parameters *γ* and *C* in the SVM used in this study gives a measure of the influence of training data points on decision boundary and a measure of miss-classification tolerance. The first parameter *γ* comes from the radial basis function kernel (e.g., 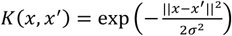 or equivalently *K*(*x, x*^′^) = exp(–γ||*x* – *x*^′^||^2^) with a parameter *γ*) where 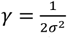. In this study, the radial basis kernel is used as a transformation function. A larger value of *γ* implies smaller *s*, which means that the classifier takes into account the effect of samples closer to the decision boundary. On the other hand, smaller *γ* means that the classifier considers the effect of samples fartherfrom the decision boundary. The *C* is a parameter of SVM that acts as regularization. It provides the classifier a trade-off between the margin of decision boundary and miss- classification. A larger value of *C* produces a narrower (smaller-margin) hyperplane if that obtains less or no miss-classification. Whereas the smaller value of *C* allows drawing a wider (bigger-margin) hyperplane even if there are some miss- classifications. The optimal value of *γ* and *C* depends on data which is why we used a grid search to tune these parameters in our classification model.

## REFERENCES

Alain, C. (2007). Breaking the wave: Effects of attention and learning on concurrent sound perception. Hearing Research, 229(1–2), 225–236.

Al-Fahad, R., Yeasin, M., & Bidelman, G. M. (2020). Decoding of single-trial EEG reveals unique states of functional brain connectivity that drive rapid speech categorization decisions. Journal of Neural Engineering, 17(1), 016045.

Alho, J., Green, B. M., May, P. J., Sams, M., Tiitinen, H., Rauschecker, J. P., & Jääskeläinen, I. P. (2016). Early-latency categorical speech sound representations in the left inferior frontal gyrus. Neuroimage, 129, 214–223.

Belin, P., McAdams, S., Thivard, L., Smith, B., Savel, S., Zilbovicius, M., Samson, S., & Samson, Y. (2002). The neuroanatomical substrate of sound duration discrimination. Neuropsychologia, 40(12), 1956–1964.

Bidelman, G. M. (2015). Induced neural beta oscillations predict categorical speech perception abilities. Brain and Language, 141, 62–69.

Bidelman, G. M., & Alain, C. (2015). Musical training orchestrates coordinated neuroplasticity in auditory brainstem and cortex to counteract age-related declines in categorical vowel perception. Journal of Neuroscience, 35(3), 1240–1249.

Bidelman, G. M., Bush, L., & Boudreaux, A. (2020a). Effects of noise on the behavioral and neural categorization of speech. Frontiers in Neuroscience, 14, 153.

Bidelman, G. M., Bush, L. C., & Boudreaux, A. M. (2020b). Effects of noise on the behavioral and neural categorization of speech. Frontiers in Neuroscience, 14, 153.

Bidelman, G. M., & Howell, M. (2016). Functional changes in inter- and intra-hemispheric cortical processing underlying degraded speech perception. NeuroImage, 124(Pt A), 581–590. https://doi.org/10.1016/j.neuroimage.2015.09.020

Bidelman, G. M., & Lee, C.-C. (2015a). Effects of language experience and stimulus context on the neural organization and categorical perception of speech. Neuroimage, 120, 191–200.

Bidelman, G. M., & Lee, C.-C. (2015b). Effects of language experience and stimulus context on the neural organization and categorical perception of speech. Neuroimage, 120, 191–200.

Bidelman, G. M., Moreno, S., & Alain, C. (2013b). Tracing the emergence of categorical speech perception in the human auditory system. Neuroimage, 79, 201–212.

Bidelman, G. M., Moreno, S., & Alain, C. (2013a). Tracing the emergence of categorical speech perception in the human auditory system. Neuroimage, 79, 201–212.

Bidelman, G. M., Moreno, S., & Alain, C. (2013c). Tracing the emergence of categorical speech perception in the human auditory system. Neuroimage, 79, 201–212.

Bidelman, G. M., & Walker, B. (2019a). Plasticity in auditory categorization is supported by differential engagement of the auditory-linguistic network. NeuroImage, 201, 116022.

Bidelman, G. M., & Walker, B. (2019b). Plasticity in auditory categorization is supported by differential engagement of the auditory-linguistic network. NeuroImage, 201, 116022.

Bidelman, G. M., & Walker, B. S. (2017a). Attentional modulation and domain-specificity underlying the neural organization of auditory categorical perception. European Journal of Neuroscience, 45(5), 690–699.

Bidelman, G. M., & Walker, B. S. (2017b). Attentional modulation and domain-specificity underlying the neural organization of auditory categorical perception. European Journal of Neuroscience, 45(5), 690–699.

Bidelman, G. M., Weiss, M. W., Moreno, S., & Alain, C. (2014). Coordinated plasticity in brainstem and auditory cortex contributes to enhanced categorical speech perception in musicians. European Journal of Neuroscience, 40(4), 2662–2673.

Binder, J. R., Liebenthal, E., Possing, E. T., Medler, D. A., & Ward, B. D. (2004). Neural correlates of sensory and decision processes in auditory object identification. Nature Neuroscience, 7(3), 295–301.

Caplan, D. (1994). Language and the brain. Academic Press, 1023–1053.

Casale, S., Russo, A., Scebba, G., & Serrano, S. (2008). Speech Emotion Classification Using Machine Learning Algorithms. 2008 IEEE International Conference on Semantic Computing, 158–165. https://doi.org/10.1109/ICSC.2008.43

Celsis, P., Doyon, B., Boulanouar, K., Pastor, J., Démonet, J.-F., & Nespoulous, J.-L. (1999). ERP correlates of phoneme perception in speech and sound contexts. Neuroreport, 10(7), 1523–1527.

Chang, E. F., Rieger, J. W., Johnson, K., Berger, M. S., Barbaro, N. M., & Knight, R. T. (2010). Categorical speech representation in human superior temporal gyrus. Nature Neuroscience, 13(11), 1428.

Cruz, J. A., & Wishart, D. S. (2006). Applications of machine learning in cancer prediction and prognosis. Cancer Informatics, 2, 117693510600200030.

de Taillez, T., Kollmeier, B., & Meyer, B. T. (2020). Machine learning for decoding listeners’ attention from electroencephalography evoked by continuous speech. European Journal of Neuroscience, 51(5), 1234–1241.

Desai, R., Liebenthal, E., Waldron, E., & Binder, J. R. (2008). Left posterior temporal regions are sensitive to auditory categorization. Journal of Cognitive Neuroscience, 20(7), 1174–1188.

Deschamps, I., Baum, S. R., & Gracco, V. L. (2014). On the role of the supramarginal gyrus in phonological processing and verbal working memory: Evidence from rTMS studies. Neuropsychologia, 53, 39–46.

Desikan, R. S., Ségonne, F., Fischl, B., Quinn, B. T., Dickerson, B. C., Blacker, D., Buckner, R. L., Dale, A. M., Maguire, R. P., Hyman, B. T., Albert, M. S., & Killiany, R. J. (2006). An automated labeling system for subdividing the human cerebral cortex on MRI scans into gyral based regions of interest. NeuroImage, 31(3), 968–980. https://doi.org/10.1016/j.neuroimage.2006.01.021

Domenech, P., & Dreher, J.-C. (2010). Decision threshold modulation in the human brain. Journal of Neuroscience, 30(43), 14305–14317.

Du, Y., Buchsbaum, B. R., Grady, C. L., & Alain, C. (2016). Increased activity in frontal motor cortex compensates impaired speech perception in older adults. Nature Communications, 7, 12241. https://doi.org/10.1038/ncomms12241

Dufor, O., Serniclaes, W., Sprenger-Charolles, L., & Démonet, J.-F. (2007). Top-down processes during auditory phoneme categorization in dyslexia: A PET study. Neuroimage, 34(4), 1692–1707.

Efron, B., Hastie, T., Johnstone, I., & Tibshirani, R. (2004). Least angle regression. The Annals of Statistics, 32(2), 407–499.

Eimas, P. D., Siqueland, E. R., Jusczyk, P., & Vigorito, J. (1971). Speech perception in infants. Science, 171(3968), 303–306.

Feng, G., Gan, Z., Wang, S., Wong, P. C., & Chandrasekaran, B. (2018). Task-general and acoustic-invariant neural representation of speech categories in the human brain. Cerebral Cortex, 28(9), 3241–3254.

Fox, R. A. (1984). Effect of lexical status on phonetic categorization. Journal of Experimental Psychology: Human Perception and Performance, 10(4), 526.

Friedman, J., Hastie, T., & Tibshirani, R. (2010). Regularization Paths for Generalized Linear Models via Coordinate Descent. Journal of Statistical Software, 33(1), 1–22.

Frost, J. A., Binder, J. R., Springer, J. A., Hammeke, T. A., Bellgowan, P. S., Rao, S. M., & Cox, R. W. (1999). Language processing is strongly left lateralized in both sexes: Evidence from functional MRI. Brain, 122(2), 199–208.

Geiser, E., Zaehle, T., Jancke, L., & Meyer, M. (2008). The neural correlate of speech rhythm as evidenced by metrical speech processing. Journal of Cognitive Neuroscience, 20(3), 541–552.

Guenther, F. H., Nieto-Castanon, A., Ghosh, S. S., & Tourville, J. A. (2004). Representation of sound categories in auditory cortical maps. Journal of Speech, Language, and Hearing Research.

Hampshire, A., Chamberlain, S. R., Monti, M. M., Duncan, J., & Owen, A. M. (2010). The role of the right inferior frontal gyrus: Inhibition and attentional control. Neuroimage, 50(3), 1313–1319.

Hanley, J. A. (1983). Appropriate uses of multivariate analysis. Annual Review of Public Health, 4(1), 155–180.

Hickok, G., Costanzo, M., Capasso, R., & Miceli, G. (2011). The role of Broca’s area in speech perception: Evidence from aphasia revisited. Brain and Language, 119(3), 214–220.

Hickok, G., Erhard, P., Kassubek, J., Helms-Tillery, A. K., Naeve-Velguth, S., Strupp, J. P., Strick, P. L., & Ugurbil, K. (2000). A functional magnetic resonance imaging study of the role of left posterior superior temporal gyrus in speech production: Implications for the explanation of conduction aphasia. Neuroscience Letters, 287(2), 156–160.

Hickok, G., & Poeppel, D. (2000). Towards a functional neuroanatomy of speech perception. Trends in Cognitive Sciences, 4(4), 131–138.

Hickok, G., & Poeppel, D. (2004). Dorsal and ventral streams: A framework for understanding aspects of the functional anatomy of language. Cognition, 92(1–2), 67–99.

Holt, L. L., & Lotto, A. J. (2010). Speech perception as categorization. Attention, Perception, & Psychophysics, 72(5), 1218–1227.

Hsu, C.-W., Chang, C.-C., & Lin, C. J. (2003). A practical guide to support vector classification technical report department of computer science and information engineering. National Taiwan University, Taipei.

Hull, R., & Vaid, J. (2006). Laterality and language experience. Laterality, 11(5), 436–464.

Husain, F. T., Fromm, S. J., Pursley, R. H., Hosey, L. A., Braun, A. R., & Horwitz, B. (2006). Neural bases of categorization of simple speech and nonspeech sounds. Human Brain Mapping, 27(8), 636–651.

James, G., Witten, D., Hastie, T., & Tibshirani, R. (2013). An introduction to statistical learning (Vol. 112). Springer.

Klingberg, T., Forssberg, H., & Westerberg, H. (2002). Increased brain activity in frontal and parietal cortex underlies the development of visuospatial working memory capacity during childhood. Journal of Cognitive Neuroscience, 14(1), 1–10.

Kuhl, P. K., Williams, K. A., Lacerda, F., Stevens, K. N., & Lindblom, B. (1992). Linguistic experience alters phonetic perception in infants by 6 months of age. Science, 255(5044), 606–608.

Lee, Y.-S., Turkeltaub, P., Granger, R., & Raizada, R. D. (2012). Categorical speech processing in Broca’s area: An fMRI study using multivariate pattern-based analysis. Journal of Neuroscience, 32(11), 3942–3948.

Liebenthal, E., Desai, R., Ellingson, M. M., Ramachandran, B., Desai, A., & Binder, J. R. (2010a). Specialization along the left superior temporal sulcus for auditory categorization. Cerebral Cortex, 20(12), 2958–2970.

Liebenthal, E., Desai, R., Ellingson, M. M., Ramachandran, B., Desai, A., & Binder, J. R. (2010b). Specialization along the left superior temporal sulcus for auditory categorization. Cerebral Cortex, 20(12), 2958–2970.

Loui, P. (2015). A dual-stream neuroanatomy of singing. Music Perception: An Interdisciplinary Journal, 32(3), 232–241.

Luck, S. J. (2005). An introduction to the event-related potential technique (pp. 45–64). Cambridge, Ma: MIT press.

Mahmud, M. S., Ahmed, F., Al-Fahad, R., Moinuddin, K. A., Yeasin, M., Alain, C., & Bidelman, G. (2020). Decoding hearing-related changes in older adults’ spatiotemporal neural processing of speech using machine learning. Frontiers in Neuroscience, 1–14.

Mankel, K., Barber, J., & Bidelman, G. M. (2020). Auditory categorical processing for speech is modulated by inherent musical listening skills. NeuroReport, 31(2), 162–166.

Martin, S., Brunner, P., Holdgraf, C., Heinze, H.-J., Crone, N. E., Rieger, J., Schalk, G., Knight, R. T., & Pasley, B. N. (2014). Decoding spectrotemporal features of overt and covert speech from the human cortex. Frontiers in Neuroengineering, 7, 14.

Masmoudi, S., Dai, D. Y., & Naceur, A. (2012). Attention, representation, and human performance: Integration of cognition, emotion, and motivation. Psychology Press.

McClelland, J. L., & Elman, J. L. (1986). The TRACE model of speech perception. Cognitive Psychology, 18(1), 1–86.

Meinshausen, N., & Bühlmann, P. (2010). Stability selection. Journal of the Royal Statistical Society: Series B (Statistical Methodology), 72(4), 417–473. https://doi.org/10.1111/j.1467-9868.2010.00740.x

Menon, V., & Desmond, J. E. (2001). Left superior parietal cortex involvement in writing: Integrating fMRI with lesion evidence. Cognitive Brain Research, 12(2), 337–340.

Miller, C. T., & Cohen, Y. E. (2010). Vocalization processing. Primate Neuroethology, 237–255.

Miller, E. K., Freedman, D. J., & Wallis, J. D. (2002). The prefrontal cortex: Categories, concepts and cognition. Philosophical Transactions of the Royal Society of London. Series B: Biological Sciences, 357(1424), 1123–1136.

Miller, E. K., Nieder, A., Freedman, D. J., & Wallis, J. D. (2003). Neural correlates of categories and concepts. Current Opinion in Neurobiology, 13(2), 198–203.

Moinuddin, K. A., Yeasin, M., & Bidelman, G. M. (2019, September 9). BrainO. https://github.com/cvpia-uofm/BrainO

Molfese, D., Key, A. P. F., Maguire, M., Dove, G. O., & Molfese, V. J. (2005). Event-related evoked potentials (ERPs) in speech perception. The Handbook of Speech Perception, 99121.

Mostert, P., Kok, P., & De Lange, F. P. (2015). Dissociating sensory from decision processes in human perceptual decision making. Scientific Reports, 5, 18253.

Myers, E. B., & Blumstein, S. E. (2008). The neural bases of the lexical effect: An fMRI investigation. Cerebral Cortex, 18(2), 278–288.

Noe, C., & Fischer-Baum, S. (2020). Early lexical influences on sublexical processing in speech perception: Evidence from electrophysiology. Cognition, 197, 104162.

Nogueira, S., Sechidis, K., & Brown, G. (2017). On the Stability of Feature Selection Algorithms. Journal of Machine Learning Research, 18, 174–1.

Norris, D., McQueen, J. M., & Cutler, A. (2000). Merging information in speech recognition: Feedback is never necessary. Behavioral and Brain Sciences, 23, 299–325.

Novick, J. M., Trueswell, J. C., & Thompson-Schill, S. L. (2010). Broca’s area and language processing: Evidence for the cognitive control connection. Language and Linguistics Compass, 4(10), 906–924.

Nyberg, L., Marklund, P., Persson, J., Cabeza, R., Forkstam, C., Petersson, K. M., & Ingvar, M. (2003). Common prefrontal activations during working memory, episodic memory, and semantic memory. Neuropsychologia, 41(3), 371–377.

Oberhuber, M., Hope, T. M. H., Seghier, M. L., Parker Jones, O., Prejawa, S., Green, D. W., & Price, C. J. (2016). Four functionally distinct regions in the left supramarginal gyrus support word processing. Cerebral Cortex, 26(11), 4212–4226.

Oldfield, R. C. (1971). The assessment and analysis of handedness: The Edinburgh inventory. Neuropsychologia, 9(1), 97–113.

Park, Y., Luo, L., Parhi, K. K., & Netoff, T. (2011). Seizure prediction with spectral power of EEG using cost-sensitive support vector machines. Epilepsia, 52(10), 1761–1770. https://doi.org/10.1111/j.1528-1167.2011.03138.x

Paus, T., Petrides, M., Evans, A. C., & Meyer, E. (1993). Role of the human anterior cingulate cortex in the control of oculomotor, manual, and speech responses: A positron emission tomography study. Journal of Neurophysiology, 70(2), 453–469.

Perlovsky, L. (2011). Language and cognition interaction neural mechanisms. Computational Intelligence and Neuroscience, 2011.

Picton, T. W., van Roon, P., Armilio, M. L., Berg, P., Ille, N., & Scherg, M. (2000). The correction of ocular artifacts: A topographic perspective. Clinical Neurophysiology, 111(1), 53–65. https://doi.org/10.1016/S1388-2457(99)00227-8

Pisoni, D. B., & Tash, J. (1974). Reaction times to comparisons within and across phonetic categories. Perception & Psychophysics, 15(2), 285–290.

Rauschecker, J. P., & Scott, S. K. (2009). Maps and streams in the auditory cortex: Nonhuman primates illuminate human speech processing. Nature Neuroscience, 12(6), 718.

Royston, P., & Sauerbrei, W. (2008). Multivariable model-building: A pragmatic approach to regression anaylsis based on fractional polynomials for modelling continuous variables (Vol. 777). John Wiley & Sons.

Ruppert, D., & Wand, M. P. (1994). Multivariate locally weighted least squares regression. The Annals of Statistics, 1346–1370.

Russ, B. E., Lee, Y.-S., & Cohen, Y. E. (2007). Neural and behavioral correlates of auditory categorization. Hearing Research, 229(1–2), 204–212.

Sabri, M., Binder, J. R., Desai, R., Medler, D. A., Leitl, M. D., & Liebenthal, E. (2008). Attentional and linguistic interactions in speech perception. Neuroimage, 39(3), 1444–1456.

Sahin, N. T., Pinker, S., Cash, S. S., Schomer, D., & Halgren, E. (2009). Sequential processing of lexical, grammatical, and phonological information within Broca’s area. Science, 326(5951), 445–449.

Saito, T., & Rehmsmeier, M. (2015). The precision-recall plot is more informative than the ROC plot when evaluating binary classifiers on imbalanced datasets. PloS One, 10(3), e0118432.

Schneiders, J., Opitz, B., Tang, H., Deng, Y., Xie, C., Li, H., & Mecklinger, A. (2012). The impact of auditory working memory training on the fronto-parietal working memory network. Frontiers in Human Neuroscience, 6, 173.

Seabold, S., & Perktold, J. (2010). Statsmodels: Econometric and statistical modeling with python. Proceedings of the 9th Python in Science Conference, 57, 61.

Tadel, F., Baillet, S., Mosher, J. C., Pantazis, D., & Leahy, R. M. (2011). Brainstorm: A user-friendly application for MEG/EEG analysis. Computational Intelligence and Neuroscience, 2011, 8.

Tankus, A., Fried, I., & Shoham, S. (2012). Structured neuronal encoding and decoding of human speech features. Nature Communications, 3(1), 1–5.

Tervaniemi, M., & Hugdahl, K. (2003). Lateralization of auditory-cortex functions. Brain Research Reviews, 43(3), 231–246.

Toscano, J. C., Anderson, N. D., Fabiani, M., Gratton, G., & Garnsey, S. M. (2018). The time-course of cortical responses to speech revealed by fast optical imaging. Brain and Language, 184, 32–42.

Tsunada, J., & Cohen, Y. E. (2014). Neural mechanisms of auditory categorization: From across brain areas to within local microcircuits. Frontiers in Neuroscience, 8, 161.

Tzourio, N., Crivello, F., Mellet, E., Nkanga-Ngila, B., & Mazoyer, B. (1998). Functional anatomy of dominance for speech comprehension in left handers vs right handers. Neuroimage, 8(1), 1–16.

Weighted Regression in SAS, R, and Python. (n.d.). Retrieved May 27, 2020, from https://jbhender.github.io/Stats506/F17/Projects/Abalone_WLS.html

Whitwell, J. L., Duffy, J. R., Strand, E. A., Xia, R., Mandrekar, J., Machulda, M. M., Senjem, M. L., Lowe, V. J., Jack Jr, C. R., & Josephs, K. A. (2013). Distinct regional anatomic and functional correlates of neurodegenerative apraxia of speech and aphasia: An MRI and FDG-PET study. Brain and Language, 125(3), 245–252.

Wood, C. C., Goff, W. R., & Day, R. S. (1971). Auditory evoked potentials during speech perception. Science, 173(4003), 1248–1251.

Xu, Y., Gandour, J. T., & Francis, A. L. (2006). Effects of language experience and stimulus complexity on the categorical perception of pitch direction. The Journal of the Acoustical Society of America, 120(2), 1063–1074.

Yin, Q.-Y., Li, J.-L., & Zhang, C.-X. (2017). Ensembling Variable Selectors by Stability Selection for the Cox Model. Computational Intelligence and Neuroscience, 2017. https://doi.org/10.1155/2017/2747431

Zatorre, R. J., Evans, A. C., Meyer, E., & Gjedde, A. (1992). Lateralization of phonetic and pitch discrimination in speech processing. Science, 256(5058), 846–849.

